# SARS-CoV-2 neutralizing camelid heavy-chain-only antibodies as powerful tools for diagnostic and therapeutic applications

**DOI:** 10.1101/2022.03.24.485614

**Authors:** A. Schlör, S. Hirschberg, G. Ben Amor, T.L. Meister, P. Arora, S. Pöhlmann, M. Hoffmann, S. Pfaender, O. Kamal Eddin, J. Kamhieh-Milz, K. Hanack

**Affiliations:** new/era/mabs GmbH, August-Bebel-Str. 89, 14482 Potsdam, Germany; Charité – Universitätsmedizin Berlin, corporate member of Freie Universität Berlin and Humboldt Universität zu Berlin, Institute of Transfusion Medicine, Robert-Koch-Platz 4, 10115 Berlin, Germany; Wimedko GmbH, Manfred-von-Richthofen-Str. 15, 12101 Berlin, Germany; Ruhr-University Bochum, Department for Molecular and Medical Virology, Universitätsstr. 150, 44801 Bochum, Germany; Infection Biology Unit, German Primate Center, Kellnerweg 4, 37077 Göttingen, Germany; Faculty of Biology and Psychology, Georg-August-University Göttingen, Wilhelmsplatz 1, 37073 Göttingen, Wimedko GmbH, Manfred-von-Richthofen-Str. 15, 12101 Berlin, Germany; University of Potsdam, Karl-Liebknecht-Str. 24-25, 14476 Potsdam, Germany

**Keywords:** camelid heavy-chain-only antibodies, single domain antibodies, nanobodies, SARS-CoV-2, neutralization, Omicron

## Abstract

**Introduction:** The ongoing COVID-19 pandemic situation caused by SARS-CoV-2 and variants of concern such as B.1.617.2 (Delta) and recently, B.1.1.529 (Omicron) is posing multiple challenges to humanity. The rapid evolution of the virus requires adaptation of diagnostic and therapeutic applications.

**Objectives:** In this study, we describe camelid heavy-chain-only antibodies (hcAb) as useful for novel *in vitro* diagnostic assays and for therapeutic applications due to their neutralizing capacity.

**Methods:** Five antibody candidates were selected out of a naïve camelid library by phage display and expressed as full-length IgG2 antibodies. The antibodies were characterized by Western blot, enzyme-linked immunosorbent assays, surface plasmon resonance with regard to their specificity to the recombinant SARS-CoV-2 Spike protein and to SARS-CoV-2 virus-like particles. Neutralization assays were performed with authentic SARS-CoV-2 and pseudotyped viruses (wildtype and Omicron).

**Results:** All antibodies efficiently detect recombinant SARS-CoV-2 Spike protein and SARS-CoV-2 virus-like particles in different ELISA setups. The best combination was shown with hcAb B10 as catcher antibody and HRP-conjugated hcAb A7.2 as the detection antibody. Further, four out of five antibodies potently neutralized authentic wildtype SARS-CoV-2 and particles pseudotyped with the SARS-CoV-2 Spike proteins of the wildtype and Omicron variant, sublineage BA.1 at concentrations between 0.1 and 0.35 ng/mL (ND50).

**Conclusion:** Collectively, we report novel camelid hcAbs suitable for diagnostics and potential therapy.

**Graphical Abstract:** 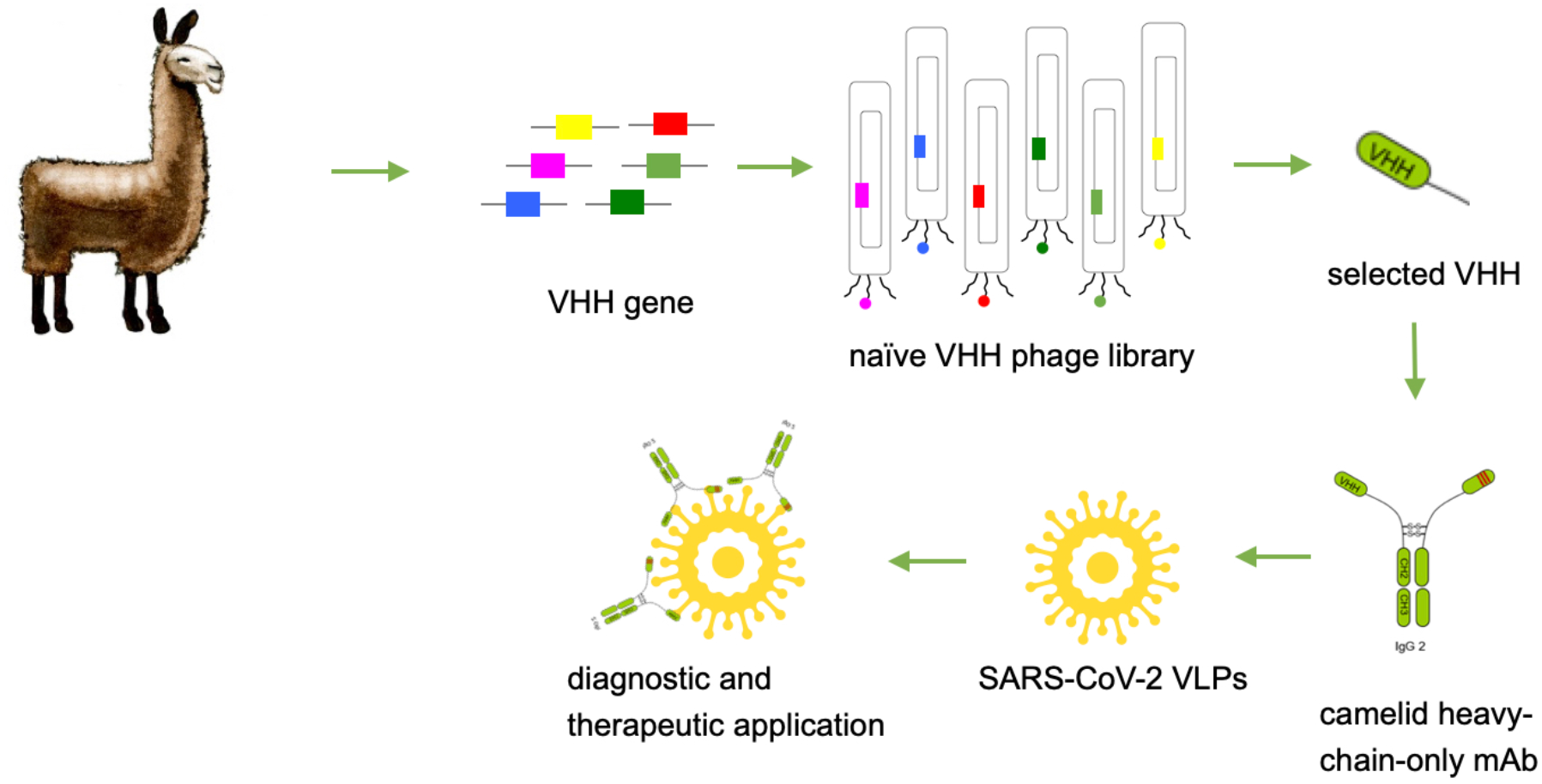

## Introduction

Camelid heavy-chain-only antibodies (hcAbs) are described as a novel class of binding molecules with highly promising advantages for diagnostic and therapeutic applications. Compared to mouse or human antibodies, hcAbs recognize their corresponding target only with one chain instead of a heavy and a light chain. The variable element of these heavy-chain-only antibodies is defined as camelid single domain antibody (VHH) or briefly as nanobody [1–2]. Nanobodies are generated by phage display from naïve or immune libraries and represent the smallest recognition unit in the class of antibody molecules with a molecular size of 15 kDa. They are characterized by a high thermal stability, a very good tissue penetration, and the recognition of hidden or difficult-to-reach epitopes [3]. Therefore, these antibodies represent a new class of next-generation antibodies and are of high interest for biomedical applications. For therapeutic use it is favorable to use the 15 kDa fragment compared to full length antibodies to avoid antibody-dependent enhancement (ADE) which is often seen for human or humanized therapeutic antibodies [4]. Further, nanobodies can be expressed recombinantly in prokaryotic expression systems which is compared to the production of full length antibodies a cheap and easy to scale up alternative [5]. The first nanobody-based drug Caplacizumab was approved recently by the FDA and EMA [6].

For *in vitro* diagnostic applications such as in ELISA systems, flow cytometry or immunofluorescence, nanobodies have to be expressed with a corresponding tag in order to provide a sufficient detection [7]. Beside different tags, it is advantageous to produce these nanobodies with a corresponding Fc part to create flexible opportunities for detection and to allow a system which is compatible with standard procedures such as biotin/streptavidin or enzyme-linked reactions.

There are already several nanobodies described which also neutralize SARS-CoV-2 [8–10]. The possibility to design multimeric formats was shown to significantly increase neutralizing effects. Nanobodies have entered the antibody field and gain more and more attention due to their unique properties. As camelid full length antibodies they represent a new opportunity for the creation of novel *in vitro* diagnostic assay formats. Here, we describe the identification and recombinant production of camelid full length antibodies from a naïve camelid library that allow detection of SARS-CoV-2 with a high sensitivity and neutralize SARS-CoV-2 with a high potency.

## Material and methods

### Library construction and VHH-biopanning

A naïve camelid VHH library was generated as previously described by Schlör et. al [11]. Briefly, RNA was isolated from peripheral blood mononuclear cells (PBMC; NucleoSpin RNA Mini Plus, Macherey-Nagel GmbH & Co.KG, Düren, Germany). For cDNA first strand synthesis RevertAid kit (Thermo Scientific) was used with a combination of random hexamer and oligo dT primers. Following a 2-step PCR protocol for VHH amplification with the primer combination P12/13 and NEM01-03_PhD. Primer sequences are listed in Table 1. Amplified VHHs were cloned into phagemid vector pADL-22c (Antibody Design Labs). The library was transformed into *E. coli* XL1 blue (Agilent) by electroporation.

**Table 1:**
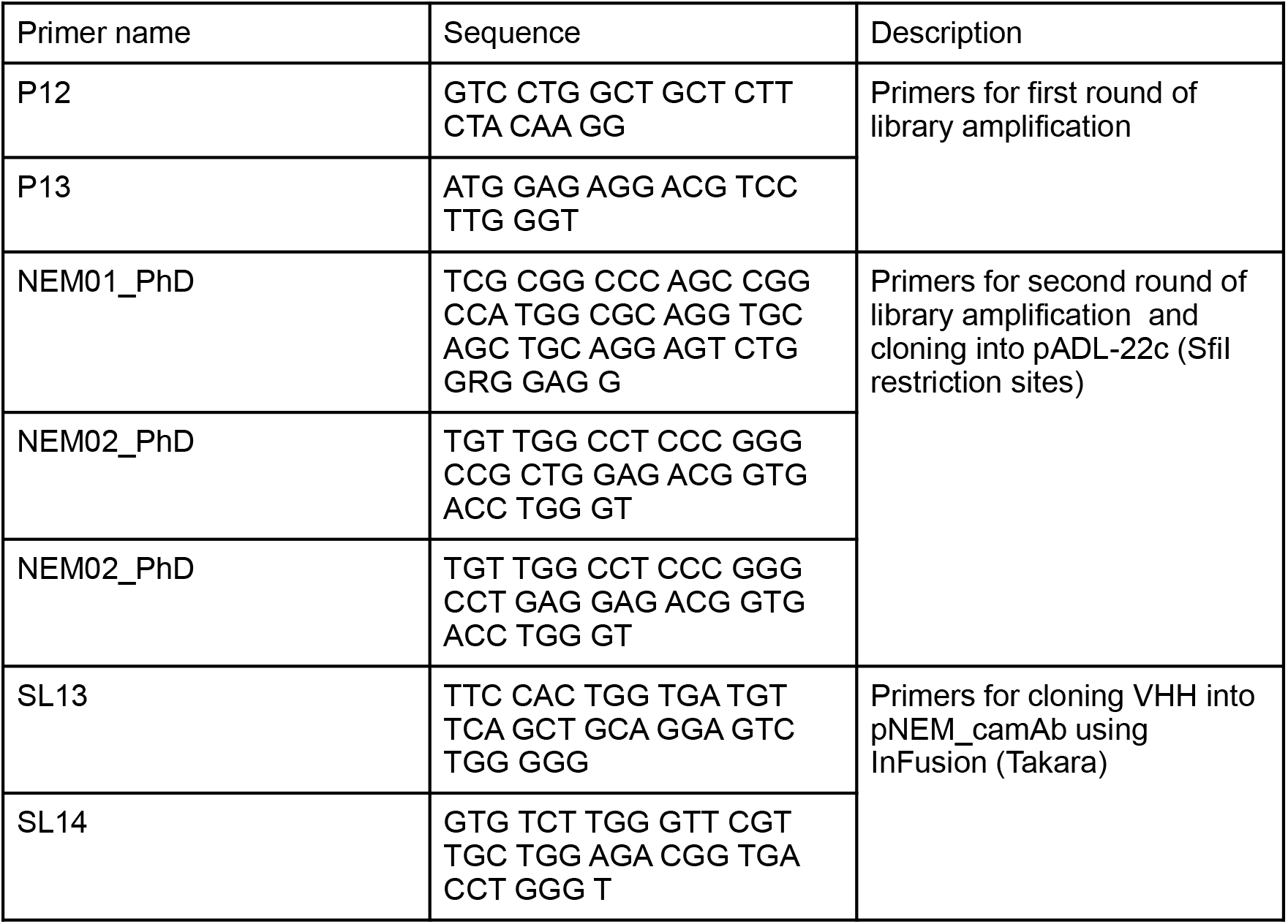
Primer sequences for amplification of VHH sequences and cloning.

For biopanning, transformed XL1 blue were cultured and inoculated with M13KO7 helper phages. The resulting phages were enriched in five rounds of biopanning. Full length SARS-CoV-2 Spike protein (SPN-C52H4; AcroBiosystems) was used as antigen in decreasing amounts starting from 15 to 0.1 μg. Post-panning analyses were performed as described by Coomber [12]. Monoclonal phages were tested in phage enzyme-linked immunosorbent assays (ELISA) for antigen specific binding to recombinant SARS-CoV-2 Spike protein. Positive clones were sequenced and become heavy chain only antibodies by cloning into the expression plasmid pNEM_camAb.

### Heavy chain Ab design, expression and purification

For the generation of a full length heavy-chain-only antibody, the different VHHs were amplified with SL13/14 and cloned into the vector pNEM_camhAb. pNEM_camAb was designed in our lab as vector for eukaryotic protein expression with a camelid Fc fragment (CH2-CH3 domain) resulting in hcAbs.

HcAbs were expressed using the Expi293 expression system (Thermo Fisher Scientific). Culturing and transfection was performed according to the manufacturer’s protocol. After 5 to 6 days after transfection, the cell culture supernatants were harvested and filtered, followed by protein A purification (ProSep^®^ Ultra Plus; Merck). Elution of hcAbs was achieved by a pH shift with glycine buffer as described previously [13]. Finally, the hcAbs were dialyzed against PBS (pH 7.4). The integrity and purity were analyzed using SDS PAGE, Western Blot and ELISA.

### Western Blot analyses and ELISA

To investigate the different hcAb candidates in Western Blot, 1 μg SARS-CoV-2 Spike protein (antibodies-online, ABIN6952734) per lane was applied onto a 4-12% SDS polyacrylamide gradient gel. The gel was blotted using the NuPage system and and Bis-Tris buffer as transfer buffer. To block unspecific binding, Rotiblock was used as blocking agent and for the dilution of the first and the second antibody. Five different hcAb candidates were applied in a concentration of 1 μg/mL (diluted in Rotiblock) and incubated for 18 h at room temperature (RT). After washing the membrane, a horseradish peroxidase (HRP) conjugated secondary antibody (ABIN1981272, antibodies-online) was applied and incubated for 3 h at RT. WesternBright Sirius was applied as substrate solution to visualize the specific signal (A44241, Invitrogen, Carlsbad, CA, USA).

For the characterization of the binding different ELISA formats were used. Indirect ELISAs were performed by coating microtiter plates with 0.5 - 1 μg/mL full length SARS-CoV-2 Spike protein (SPN-C52H4; AcroBiosystems) in 1x PBS overnight at 4 °C (50 μL/ well), various hcAb concentrations (50 μL/well in PBS/1% casein or PBS/5% neonatal calf serum (NCS), 1 h incubation at RT) and 1-2 μg/mL of HRP-conjugated secondary antibody (ABIN1981272, antibodies-online) in 50 μL/well supplemented with PBS/1% casein or PBS/5 % NCS for 45 min incubation at RT.

For the phage ELISA, the wells were coated with full length SARS-CoV-2 Spike protein (3 μg/mL in PBS, 50 μL/well) and incubated with monophages (50 μL/well). Detection of the bound phages was done with HRP-conjugated anti-M13 (clone B62, produced in our lab) in a dilution of 1:8000. Sandwich ELISAs were performed with hcAbs in different concentrations on coated antigen as described above. Each ELISA was finished using TMB as substrate solution (50 μL/well, Carl Roth) and 1 M H_2_SO_4_ as stop solution. To block nonspecific protein binding, 1 % casein in PBS or 5 % NCS in PBS was used. Between the single steps the wells were washed 3 times with tap water and 10 times after the secondary antibody incubation. Optical density was measured at 450 nm with a 620 nm reference.

### HRP conjugation of hcAbs B10 and A7.2

HRP-conjugation of hcAbs was done using a modified version of the classical periodate method as published in 1985 [14]. For activation of HRP, 0.5 mg/mL HRP in 100 mM NaHCO_3_ (pH 8.1) were mixed 1:1 with 12.5 mM NaIO_4_ and incubated at RT for 2 h in the dark. In the next step, the same volume of activated HRP and hcAbs (1 mg/mL in NaHCO_3_; pH 9.2) were incubated in a glas-wool plugged Pasteur pipet filled with Sephadex G-25 (GE Healthcare) for 3 h at RT in the dark. The elution of conjugated hcAbs from Sephadex was performed with 100 mM NaHCO_3_ (pH 9.2). To stop the reaction, 1/20th volume of 5 mg/mL NaBH_4_ was added and incubated at 4 °C. After 30 min another volume (1/10) of freshly prepared NaBH4 solution was added and incubated at 4 °C for 1 h. Finally, after an ammonium sulfate precipitation overnight HRP conjugated hcAbs were dissolved in PBS (pH 7.4).

### SPR analysis

The hcAb-binding properties of B10 were analyzed at 25 °C on a Biacore T200 instrument (GE Healthcare) using 10 mM HEPES pH 7.4, 300 mM NaCl, 3 mM EDTA, 0.05% Tween 20, 0.25 mg/ml BSA as running buffer. SARS-CoV-2 Spike protein-RBD-mFc (Acrobiosystem; SPD-C5259) was captured by a covalently immobilized antimouse IgG on a C1 sensor chip. Increasing concentrations of hcAb B10 (0.23 - 60 nM) were injected. Analyte responses were corrected for unspecific binding and buffer responses. The sensorgrams was fitted with a 1:1 binding model using Biacore evaluation software.

### Neutralization Assay

To determine the neutralization capacity of hcAbs against SARS-CoV-2, a neutralization assay with a propagation-incompetent VSV*ΔG pseudovirus system or full length virus was performed as previously described [15]. The expression vector for the Omicron spike (based on isolate hCoV19/Botswana/R40B58_BHP_3321001245/2021; GISAID Accession ID: EPI_ISL_6640919) was generated by Gibson assembly using five overlapping DNA strings (Thermo Fisher Scientific, sequences available upon request), linearized (BamHI/XbaI digest) pCG1 plasmid and GeneArt™ Gibson Assembly HiFi Master Mix (Thermo Fisher Scientific). Gibson assembly was performed according to manufacturer’s instructions. The pCG1 vector was kindly provided by Roberto Cattaneo (Mayo Clinic College of Medicine, Rochester, MN, USA). In specific, VSV*ΔG either bearing the SARS-CoV-2 (D614G) or SARS-CoV-2 B.1.1.529 (Omicron) Spike protein was incubated with a two-fold dilution of hcAbs and subsequently used to inoculated VeroE6 cells. Firefly luciferase (FLuc) activity was determined 18 hours post infection, and the reciprocal antibody dilution causing 50% inhibition of the luciferase reporter was calculated. In a similar experimental setup full length WT virus hCoV-19/Germany/BY-Bochum-1/2020 (B.1.1.70) (GISAID accession ID: EPI_ISL_1118929) [16] was incubated with a serial dilution of hcAbs and hereafter transferred onto VeroE6 cells. The cells were incubated for 72 h before they were stained with crystal violet to visualize cytopathic effects.

### Generation of SARS-CoV-2 virus-like particles (VLPs)

Human codon optimized sequences of genes encoding the S and E structural proteins of SARS-CoV-2 were synthesized by BioCat GmbH (Heidelberg, Germany) and subcloned into the pcDNA3.1 expression plasmid using the NheI 5’ and XhoI 3’ restriction site, respectively. The SARS-CoV-2 Spike protein contained the D614G mutation and the furin-cleavage site was destroyed (FKO) by R683A and R685A substitution. Another plasmid (pEXP-M+N) for the dual expression of the human codon optimized sequences of the M and N protein was generated by Vectorbuilder Inc. (VB200528-1033wpt). The two proteins were expressed from the same open reading frame separated by a T2A self-cleaving peptide and controlled by the human eukaryotic translation elongation factor 1 α1 promoter (EF1A). All plasmids were transformed into high efficiency chemically competent cells DH5α (New England BioLabs, Ipswich, MA, USA) using the heat shock method in a water bath at 42°C for 30 seconds, followed by shaking incubation in SOC outgrowth media at 37°C for 45 min. Next, 50 μL of the cell-containing media were plated on LB plates containing 50 μg/mL ampicillin. After incubation at 37°C overnight, resistant single colonies were picked and amplified in LB medium. The correct constitution of the plasmid was examined by restriction enzyme digest and the open reading frames were verified by DNA sequencing (Eurofins Genomics Germany GmbH, Ebersberg, Germany). Endotoxin free DNA was prepared using respective kits (Macherey-Nagel GmbH & Co. KG, Düren, Germany), quantified with a NanoDrop Spectrophotometer (Thermo Fisher Scientific, Wilmington, DE, USA), controlled by restriction digest and subsequently transfected into mammalian cells for VLP production. Expi293™ Expression System Kit, composed of Expi293 suspension adapted cell line, Expi293 transfection reagents and Expi293 culture medium was purchased from Thermo Fisher Scientific. Expi293 cells were grown at 37°C, 8% CO2 with 130-150 rpm on a Rotamax120 platform shaker (Heidolph Instruments GmbH & Co. KG, Schwabach, Germany) in Expi293 medium containing a final concentration of 100 U/mL penicillin-streptomycin. Cell diameter, the percentage of viable cells (vitality) and the concentration of viable cells were routinely monitored using LUNA Cell Counter (Logos Biosystems, Anyang, South Korea). Cells were seeded at a density between 0.3×10^6 and 0.5×10^6 cells/mL and subcultured when concentration reached 3×10^6 to 5×10^6 viable cells/mL which was typically after 3-5 days. For transfection, cells were precipitated for 5 minutes at 300 x g and subsequently resuspended in fresh media to a final concentration of approximately 3×10^6 viable cells/mL and a vitality greater than 95%. Per 10^6 cells approximately 1 μg of total DNA was transfected. The DNA mix was composed of pcDNA3.1-Spike (FKO), pcDNA3.1-E-Protein and pEXP-M+N-protein at a ratio of 6:2:3 diluted in Opti-MEM I Medium. To generate control VLPs without the Spike protein, the plasmid containing the Spike protein gene was replaced with a mock plasmid. ExpiFectamin 293 Reagent was also diluted in Opti-MEM I medium at the optimized ratio of 3.2 μL per 1 μg of DNA before it was slowly added to the DNA mixture. The ExpiFectamin DNA complex was incubated for 15 min at RT prior to the dropwise addition onto the gently agitated cells. Enhancers 1 and 2 were added approximately 20 h after transfection according to the manufactures protocol. Next, 96-120 h after transfection when the vitality usually decreased to 40-60%, cell culture supernatants where cleared by centrifugation at 2000 x g for 10 min followed by filtration with a 1.2 μm Minisart NML (Sartorius Stedim Biotech GmbH, Göttingen, Germany) and a 0.45 μm Millex Low Binding Durapore (PVDF) syringefilter (Merk Millipore Ltd., Tullagreen, IE). VLPs were precipitated from the clarified supernatant by the addition of PEG-it Virus Precipitation Solution (System Biosciences, Palo Alto, CA, USA) at a ratio of 1:10. The supernatants were incubated at 4°C on a rotating shaker for 24-48 h prior to precipitation at 1500 x g for 30 min. The supernatant was carefully removed, and the VLP containing pellet was resuspended 0.05 % - 0.1 % of the initial volume with sterile PBS (pH 7.2). The resuspended pellets were kept at 4°C for short-term storage (1-3 weeks) or at −80°C for long-term storage.

### Nanoparticle tracking analysis (NTA)

NTA was used to measure size and concentration of VLPs in different preparations. NTA measurements were performed using a NanoSight LM10 instrument (NanoSight, Amesbury, UK) consisting of a conventional optical microscope, high sensitivity sCMOS camera and a LM10 unit equipped with a 488 nm laser module. The samples were injected into the LM unit via the nanosight syringe pump with a constant flow rate of 50 μL/min with a 1 mL sterile syringe. Samples were diluted 1:5000. The capturing settings (shutter and gain) and analyzing settings were manually adjusted and kept constant between all samples that were recorded on the same day. NTA software (NTA 3.2 Dev Build 3.2.16) was used to capture three videos of 30 seconds and to analyze nanoparticle tracking data per sample.

## Results

### Development of hcAbs

Camelid single domain antibodies were selected by phage display out of a naïve camelid library as previously described [11]. We could identify around 15 monophages with a specific ELISA signal of more than 0.5 when recombinant full length SARS-CoV-2 Spike protein was used for coating (Fig. 1). After sequencing, we chose 5 different candidates - A7.2, B10, D3, D12, G10 - for recombinant expression of camelid full length IgG2 antibodies. Antibodies were purified by protein A chromatography [12] and characterized by Western blot analyses for their specific recognition (Fig. 2A). All hcAbs specifically recognize the recombinant Spike protein in Western blot, whereas A7.2 and B10 showed a more intense signal than the other candidates.

**Figure 1.**
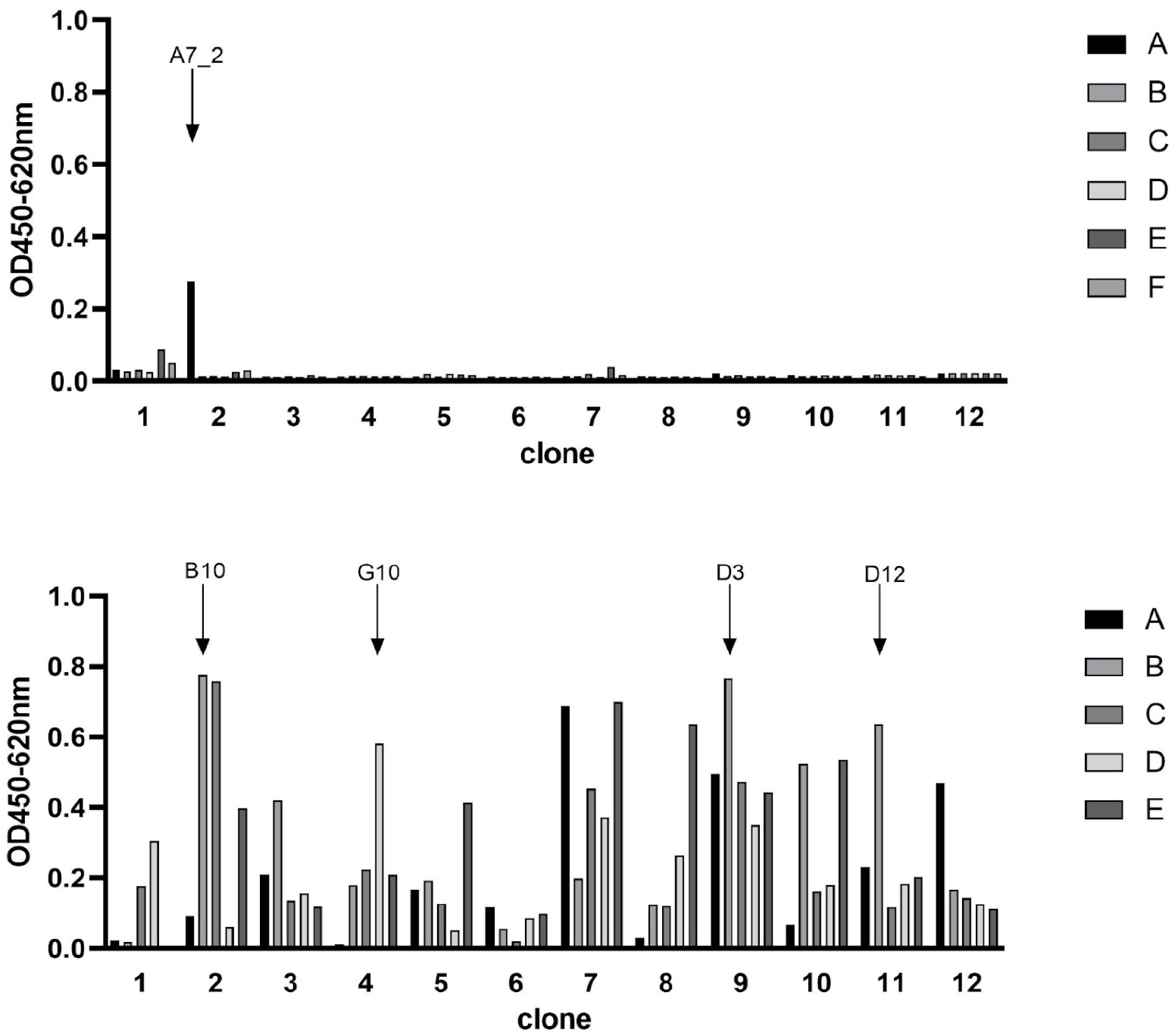
Phage display ELISA for the selection of SARS-CoV-2 Spike protein specific VHHs. Wells were coated with antigen (SARS-CoV-2 spikeprotein, 3 μg/mL) and blocked with 100 μL/well PBS/ 1% casein. Detection of specific phages was performed with a HRP-conjugated M13 antibody (diluted 1:8000). Optical density was measured at 450 nm with a reference of 620 nm. Selected candidates are labeled with arrows.

**Figure 2.**
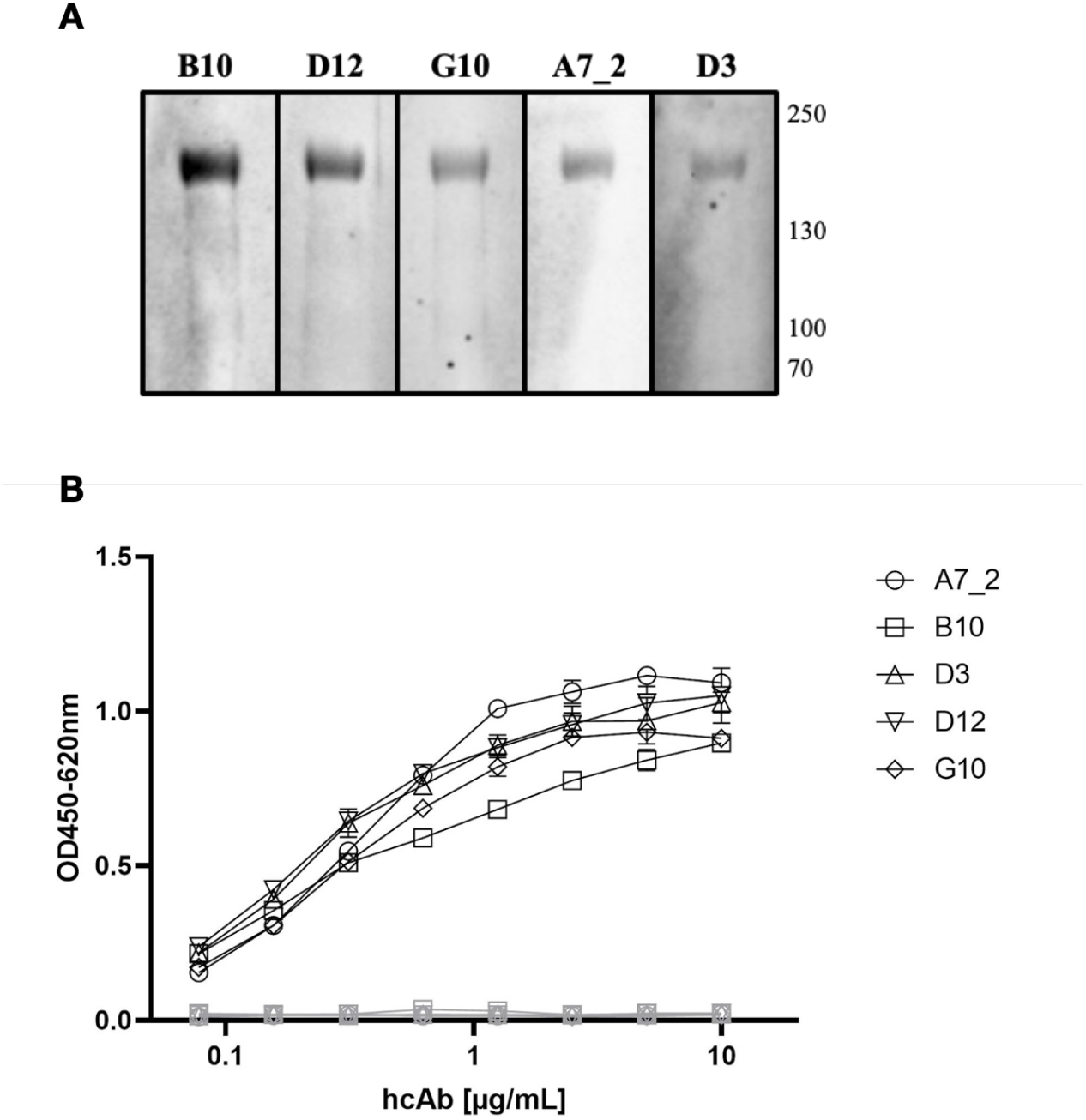
Characterization of SARS-CoV-2 Spike protein specific VHHs. **A) Western Blot analyses** Recombinant Spike protein was applied to a 4-12% SDS polyacrlyamide gradient gel and blotted using the NuPage system. HcAb candidates (1 μg/mL) were incubated for 18 h. Detection took place using a HRP-conjugated secondary antibody and WesternBright Sirius as substrate solution. **B) Indirect ELISA with recombinant SARS-CoV-2 Spike protein as antigen** Wells were coated with antigen (SARS-CoV-2 Spike protein, 3 μg/mL) and blocked with 100 μL/well PBS/ 1% casein. HcAbs were added in different dilutions. Detection was performed with a HRP-conjugated secondary antibody (ABIN1981272, 1 μg/mL) and optical density was measured at 450 nm with a reference wavelength of 620 nm. Mean and standard deviation of three independent measurements is shown.

### Characterization of hcAbs in different ELISA setups and SPR

The purified hcAbs were further characterized in different ELISA setups, to verify the optimal combinations and concentrations. First, the antibodies were serially diluted on recombinant SARS-CoV-2 Spike protein (Fig. 2B) where they showed a specific recognition till a working concentration of 0.5 μg/mL. All hcAbs showed a similar performance in this assay format.

In the next step, we tested pairwise combinations to find the optimal catcher and detector antibodies for a sandwich immunoassay principle (Fig. 3). To ensure a direct detection, we conjugated hcAb B10 and A7.2 to HRP and tested their performance in a direct ELISA (Fig. 3A). A7.2 showed a more succinct labeling with HRP and a better performance in the assay compared to B10. Therefore, we used HRP-conjugated hcAb A7.2 in a dilution of 1:1600 as detection antibody. Figure 3B shows the results for the sandwich immunoassay. We coated all hcAb candidates in different concentrations of 10, 5 and 2.5 μg/mL on the solid phase. Recombinant SARS-CoV-2 Spike protein was used as antigen and HRP-conjugated hcAb A7.2 as detection antibody. The usage of B10 as catcher hcAb and HRP-conjugated hcAb A7.2 as detector seemed to be the most sensitive combination in this sandwich immunoassay format (Fig. 3B, 2nd).

**Figure 3.**
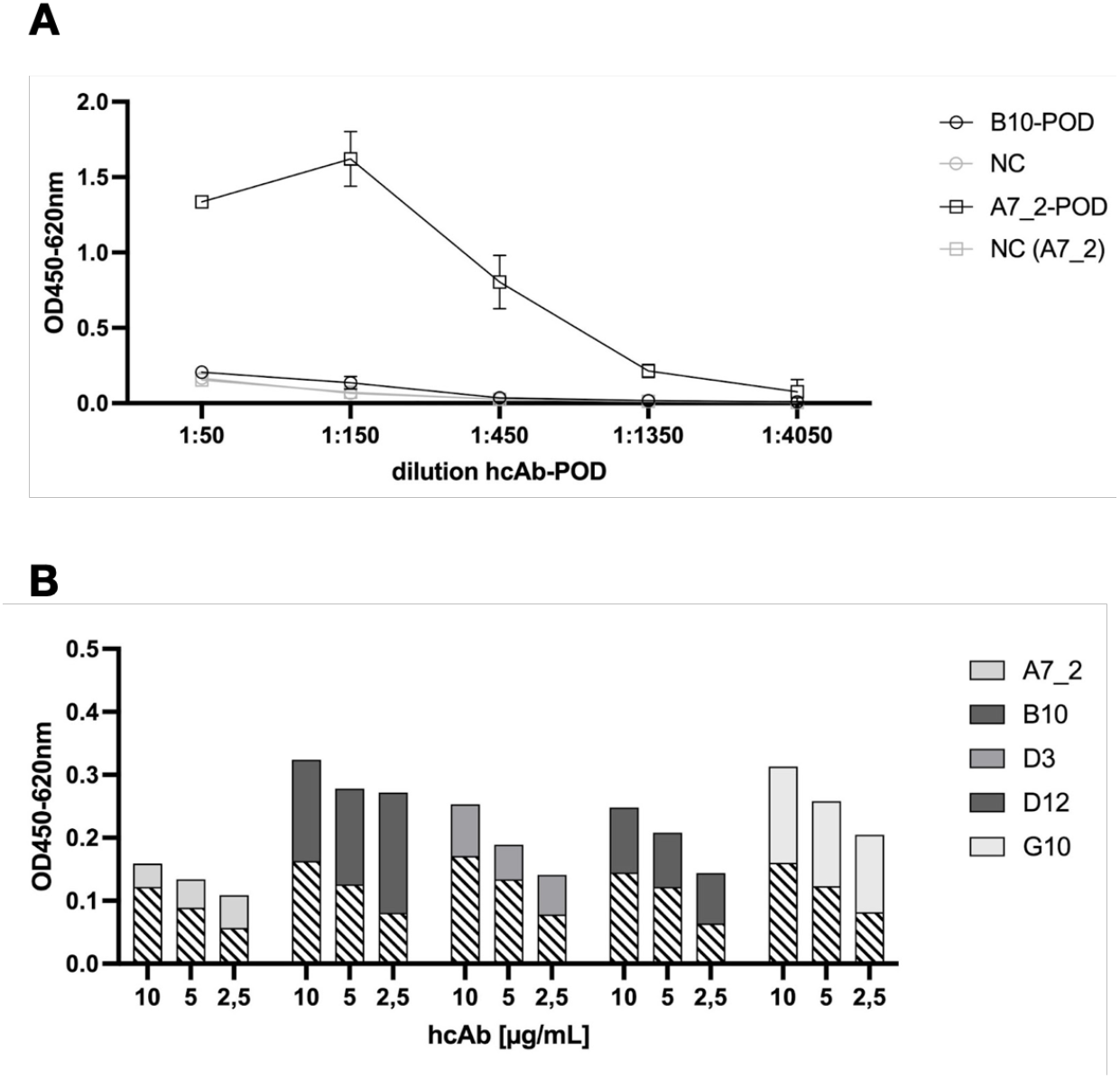
Sandwich Immunoassay setup for the detection of recombinant SARS-CoV-2 Spike protein. **A) HRP-conjugation of hcAbs B10 and A7.2** HcAbs B10 and A7.2 were conjugated with HRP to serve as secondary antibody in a sandwich ELISA. To detect the HRP conjugation, SARS-CoV-2 Spike protein was coated on the solid phase (1 μg/mL). HRP-conjugated hcAb B10 and A7.2 were added in different dilutions. Binding was detected by adding TMB substrate solution and optical density was measured at 450 nm with a reference wavelength of 620 nm. Mean and standard deviation of two independent measurements is shown. **B) Sandwich ELISA of hcAb candidates in different pairwise combinations** Purified hcAbs were coated on the solid phase in different concentrations of 10, 5 and 2.5 μg/mL. SARS-CoV-2 Spike protein was added (2.5 μg/mL). HcAb A7.2 was applied as HRP-conjugated secondary antibody in a dilution of 1:1600. Detection of optical density was performed at 450 nm with a reference wavelength of 620 nm. Cross-hatched areas represent the value for the unspecific background.

In Figure 4 we have summarized the results for our generated SARS-CoV-2 VLPs.We could prove that the generated VLPs have the expected particle size (Fig. 4A) and carry the Spike (S) as well as the nucleoprotein (N) which were detectable with human reconvalescent patient sera in Western blot experiments (Fig. 4B). The presence of E and M protein was not specifically tested because their presence is necessary for the release of VLPs from the host cell [17–18]. The VLPs were used as antigen coated on the solid phase and detected by either hcAb B10 or A7.2 followed by a HRP-conjugated secondary antibody (Fig. 4C). The dilution series revealed a working concentration of 1.25 μg/mL for hcAb B10 and 0.6 μg/mL for hcAb A7.2.

**Figure 4.**
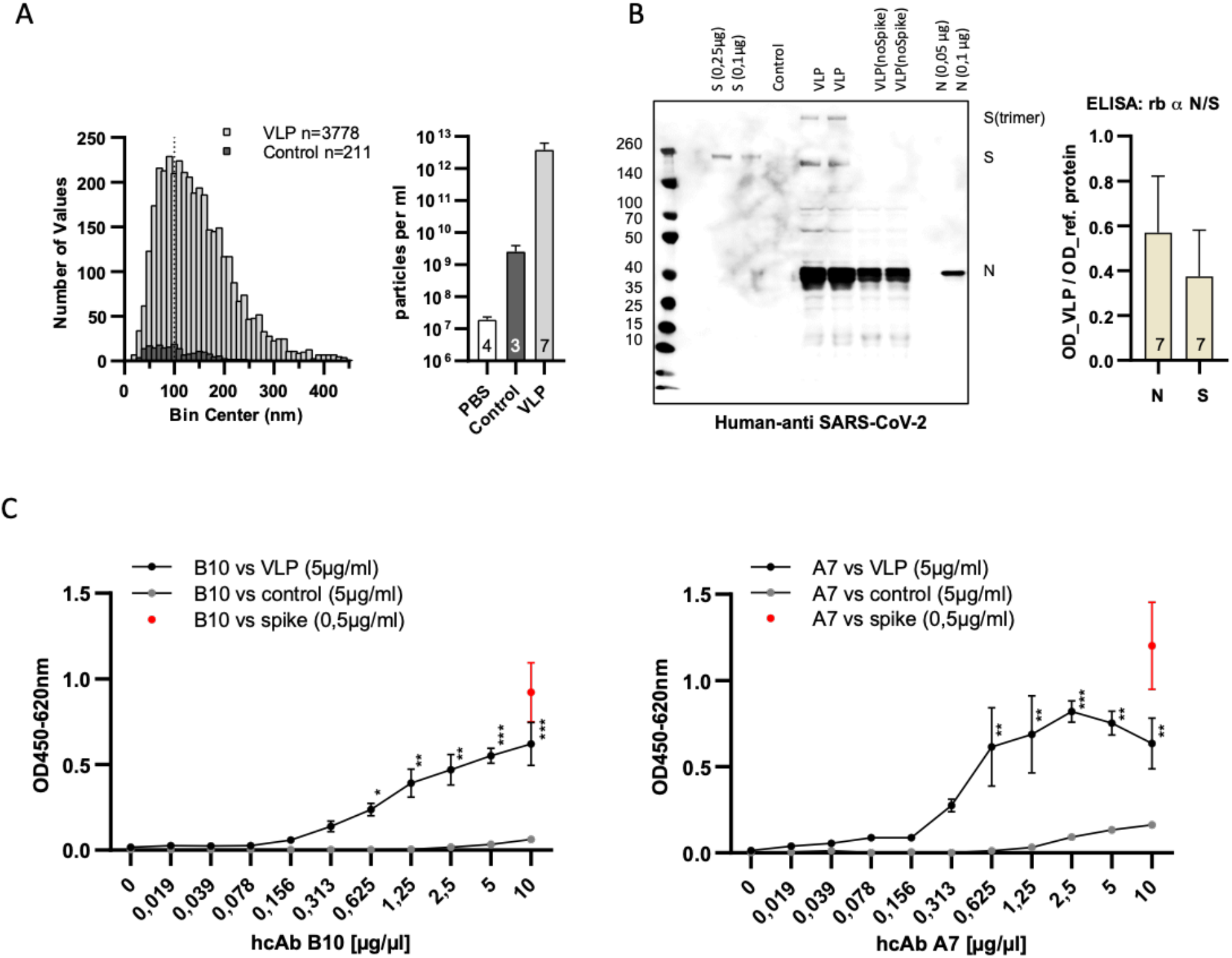
Characterization and specific binding of SARS-CoV-2 VLPs. **A) Nanoparticle tracking analysis of particle size and concentration** VLPs were purified from cell culture supernatants transfected with expression plasmids of four SARS-CoV-2 structural proteins (S, N, M and E). Data are derived from seven independent transfections (VLPs) or three untransfected cell culture supernatants (control). Samples were diluted 1:5000 prior to measuring in NanoSight LM10. The size distribution of individual particles and the particle concentration of the samples is shown. The dotted line at 100 nm indicates the approximate size of authentic SARS-CoV-2 VLPs [25]. **B) Confirmation of the presence of Spike- and nucleoprotein in SARS-CoV-2 VLPs** For the detection of Spike- and nucleoprotein on SARS-CoV-2 VLPs human reconvalescent serum was used in a dilution of 1:100 in western blot anlayses. Reference proteins served as controls. For the ELISA, VLPs (5 μg/mL) were coated on the solid phase and detected with commercial antibodies specific for the Spike- and the nucleoprotein. Reference proteins were used in a concentration of 0.25 μg/mL to serve as control. Mean values and standard deviation of seven independent VLP preparations is shown. **C) Dose-dependent detection of SARS-CoV-2 VLPs using hcAbs B10 and A7.2** VLPs, control particles or Spike protein were coated on the solid phase. HcAbs B10 and A7.2 were applied in different dilutions. Detection was performed using a HRP-conjugated secondary antibody. The optical density was measured at 450 nm with a reference wavelength of 620 nm. Mean and standard deviation of three independent measurements is shown. ELISA: 2-log dilution of B10 and A7.2. Kruskal Wallis test with Benjamini, Krieger and Yekuteli correction for multiple comparison against 0 μg/ml *** p < .001, ** p < .01, * P < .05. n=3

Next, we characterized the hcAb candidate B10 in surface plasmon resonance spectrometry (SPR). SPR measurements were performed with immobilized SARS-CoV-2 Spike protein (RBD domain) recombinantly expressed with a murine Fc part. For hcAb B10 highly reproducible kinetic data could be received, revealing a KD value of 0.39 nM (Fig. 5).

**Figure 5.**
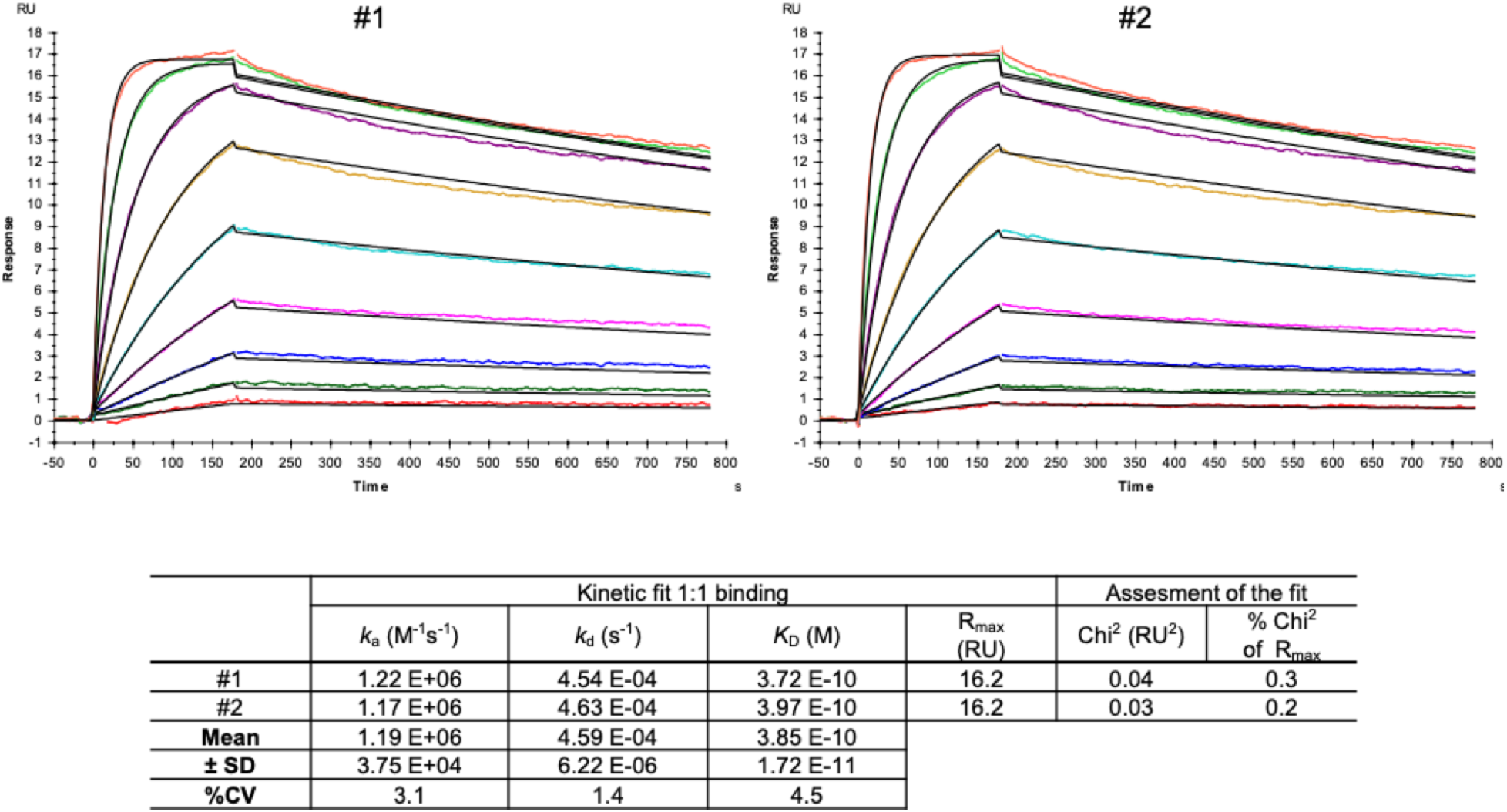
SPR measurements of hcAb B10. SARS-CoV-2 Spike protein-RBD-mFc (Acrobiosystem; SPD-C5259) was used as antigen and captured by a covalently immobilized anti-IgG1 antibody on a C1 sensor chip. Increasing concentrations of hcAb B10 (0.23 - 60 nM) were injected. Analyte responses were corrected for unspecific binding and buffer responses. The sensograms was fitted with a 1:1 binding model using Biacore evaluation software.

### Neutralization capacity of hcAbs

Neutralization assays were performed with full length SARS-CoV-2 wildtype and SARS-CoV-2 pseudotyped viruses for the wildtype, and the Omicron variant, respectively. Figure 6 shows the results for the neutralization capacity of hcAb B10, D3, and D12 for the wildtype virus (A-C), and the pseudotyped viruses of wildtype (D-F) and Omicron (G-I). B10 and D12 were strongly neutralizing the full length wildtype virus at a concentration of 1 μg/mL. D3 showed an intermediate neutralizing activity between 4.3 and 5.8 μg/mL. In neutralization assays with the pseudotype particles (Fig. 6, D-I) the hcAbs showed a similar capacity as shown for the full length wildtype virus. Surprisingly, we found that also the new variant of concern B1.1.529 (Omicron) was neutralized by the hcAbs B10, D3 and D12 in a range of 0.1 - 0.35 ng/mL (ND50). For hcAb A7.2 there was no neutralization detected (data not shown).

**Figure 6.**
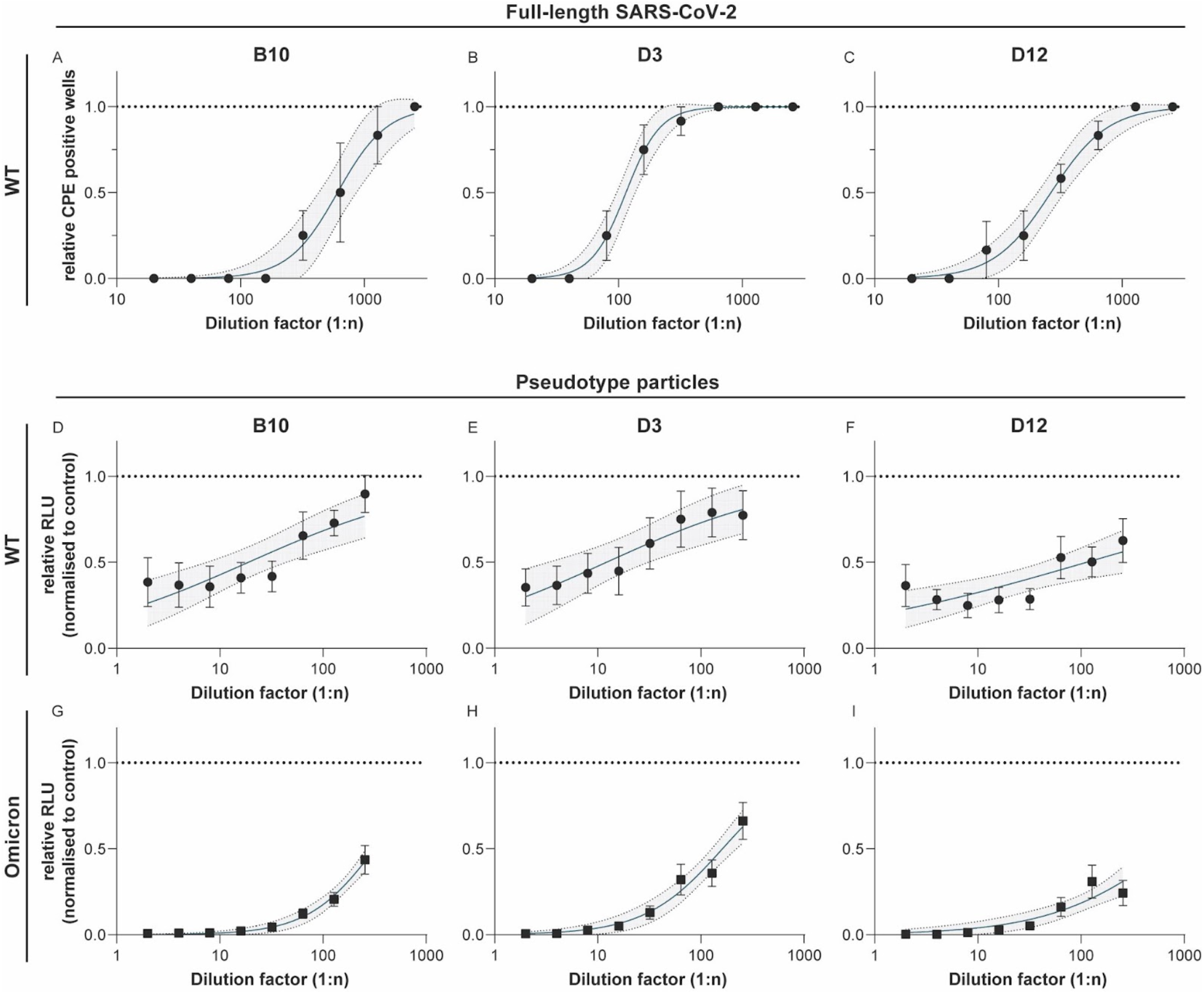
Neutralization experiments with SARS-CoV-2 wildtype virus and pseudotyped viruses (wildtype and Omicron). HcAbs B10, D3 and D13 were used for neutralization experiments with the wildtype virus hCoV-19/ Germany/BY-Bochum-1/2020 (B.1.1.70) (A-C) and with a propagation-incompetent VSV*ΔG pseudovirus system for the wildtype (D-F) and Omicron (G-H). VSV*ΔG were incubated with a two-fold dilution of hcAbs and used for infection of VeroE6 cells. Neutralization was measured using a Firefly luciferase (FLuc) reporter system. The reciprocal antibody dilution causing 50% inhibition of the luciferase reporter was calculated (ND50) in 3 independent experiments. Mean and standard deviation are shown as well as a 95% confidence interval set off with grey.

## Discussion

Camelid antibodies represent a promising antibody format for therapeutic as well as *in vitro* diagnostic applications. These types of antibodies provide favourable properties such as a high thermostability, easy and reproducible expression in prokaryotic and eukaryotic expression systems. With 15 kDa in size, nanobodies serve as the smallest known antigen binding unit and are used mainly for therapeutic applications [19]. For *in vitro* diagnostic assay systems nanobodies are not practicable due to the absent Fc chain which is needed to build up a sensitive detection. Current *in vitro* diagnostic systems use polyclonal or monoclonal full length antibodies as detector antibodies coupled to HRP or biotin to provide a measurable and sensitive detection of the target. So far, only few assay systems based on camelid nanobodies are published [20–22] but none of these are using camelid full length antibodies for detection. This study identifies suitable camelid antibody candidates specific for SARS-CoV-2 Spike protein selected by phage display from a camelid naïve library. We have expressed the nanobodies as full length heavy-chain-only antibody formats of an IgG2 isotype to establish tools for an ELISA based detection system. Normally, we would expect lower affinities when selecting candidates from a naïve library because the immune system was not specifically challenged against the target [23]. But the five candidates chosen from the selection rounds in this study showed good binding properties against the target. The hcAb B10 was characterized further with an affinity constant of 0.39 nM which is compared to other SARS-CoV-2 nanobodies from immune or synthetic libraries, a highly affine binder. This very good performance was also mirrored by the usability of hcAb B10 as catcher antibody in the developed ELISA format. Due to the high specificity it was possible to reduce the coating concentration to less than 1 μg/mL. Together with hcAb A7.2 as a detector antibody, a very potent antibody pair could be established. The successful conjugation of these hcAb candidates with routinely used enzymes such as HRP offers a standardized use of hcAbs also for *in vitro* diagnostic applications. Until now, there is no similar assay system described in literature. Our results prove the feasibility of assay systems completely designed and coordinated with camelid full length heavy-chain-only antibodies. These opportunities offer new opportunities and provide flexible tools for future applications.

Further, we could show that these hcAb candidates are not only useful as in *vitro* diagnostic tests components. The neutralization data for the full length wildtype virus as well as the pseudotyped wildtype and the new variant of concern B1.1.529 (Omicron) reveal a possible therapeutic application too. With four out of five candidates which are able to effectively neutralize the SARS-CoV-2 wildtype virus as well as the newly emerged Omicron variant we can provide highly efficient tools useful for a range of different applications. As recently described, the therapeutic potential of current human antibodies against the newly emerged Omicron variant is reduced caused by the increased number of mutations in the RBD domain and the resulting loss of specificity [24]. Our hcAb portfolio can therefore provide an alternative and improve the current availability of antibodies for both, a diagnostic as well as a therapeutic use.

## Conclusion

Camelid antibodies have a high potential for diagnostic and therapeutic applications because of their exclusive properties compared to routinely used human or mouse antibodies. The approval of the first camelid therapeutic antibody is an important step. Also for *in vitro* diagnostic assay systems camelid antibodies provide substantial advantages. They can easily be adapted to a routine detection by using corresponding Fc parts and specific secondary antibodies. To create novel tools and establish applications based on camelid antibodies will significantly impact future developments in the antibody field.

## Acknowledgements

We would like to thank Dr. Michael Zenn from Biaffin GmbH & Co.KG for performing the SPR measurements of hcAb B10. In addition we would like to thank Elena Vidal Blanco of the Department for Medical and Molecular Virology for technical support. We thank Gert Zimmer, Institute for Virology und Immunology, Switzerland and Department of Infectious Diseases and Pathobiology (DIP), Vetsuisse Faculty, University of Bern, Switzerland for providing plasmids and reagents.

## Authorship contributions

AS designed the phage display experiments and performed the panning rounds, hit selection, recombinant expression, purification and ELISA characterizations of the hcAbs. SH developed the protocols for VLP expression and characterization and preformed the Western Blot experiments. GBA contributed to the production and characterization of VLPs. TLM, PA, MH performed the neutralization experiments with authentic SARS-CoV-2 and pseudotyped viruses. SP and SP coordinated the neutralization experiments and provided the Omicron plasmid. OKE, JKM and KH initiated the project, coordinated and supervised the experiments. All authors contributed to the writing of the manuscript and agreed to the final version of it.

